# Unraveling the response of a biomimetic actin cortex to electric pulses in vesicles

**DOI:** 10.1101/338566

**Authors:** Dayinta L. Perrier, Afshin Vahid, Vaishnavi Kathavi, Lotte Stam, Lea Rems, Yuval Mulla, Aswin Muralidharan, Gijsje H. Koenderink, Michiel T. Kreutzer, Pouyan E. Boukany

**Affiliations:** Department of Chemical Engineering, Delft University of Technology, the Netherlands; AMOLF, Department of Living Matter, the Netherlands

## Abstract

We study the role of a biomimetic actin cortex during the application of electric pulses that induce electroporation or electropermeabilization, using giant unilamellar vesicles (GUVs) as a model system. The actin cortex, a subjacently attached interconnected network of actin filaments, regulates the shape and mechanical properties of the plasma membrane of mammalian cells, and is a major factor influencing the mechanical response of the cell to external physical cues. We demonstrate that the presence of an actin shell inhibits the formation of macropores in the electroporated GUVs. Additionally, experiments on the uptake of dye molecules after electroporation show that the actin network slows down the resealing process of the permeabilized membrane. We further analyze the stability of the actin network inside the GUVs exposed to high electric pulses. We find disruption of the actin layer that is likely due to the electrophoretic forces acting on the actin filaments during the permeabilization of the GUVs. Our findings on the GUVs containing a biomimetic cortex provide a step towards understanding the discrepancies between the electroporation mechanism of a living cell and its simplified model of the empty GUV.

## Introduction

The plasma membrane is a selective barrier that separates the intracellular environment from the cell exterior and regulates the influx and efflux of differently sized molecules. The membrane consists of a lipid bilayer with numerous anchored and embedded inclusions, providing the membrane with both fluid and elastic properties; additionally, an inter-connected network of actin filaments, called the actin cortex, lies immediately underneath the plasma membrane. The bi-directional interplay between the membrane and actin cortex strongly influences the mechanical response of the cell to external stimuli during diverse biological processes, ranging from cell migration and differentiation to cell division^1^.

The actin cortex also plays an essential role in electroporation, also referred to as electropermeabilization, of cells. This is a membrane permeabilization technique used for the delivery of a wide variety of molecules ranging from small molecules such as drugs to large molecules, such as DNA, into the cell. Applying direct current (DC) electric pulses to a cell builds up an induced transmembrane voltage, which can permeabilize the cell membrane above a critical transmembrane voltage (∼0.2 - 1 V)^2, 3^. The kinetics of electroporation in cells involves five consecutive steps: (i) nucleation of defects in the membrane, (ii) expansion of the defects, (iii) stabilization of the permeabilized state, (iv) resealing of the permeated membrane, (v) “memory effect” including a full structural recovery of the already resealed membrane^3^. It has been shown that cells can remain permeabilized from minutes up to hours after pulsation^4–6^. This long resealing process is considerably slower than that observed for bare lipid bilayers (less than one second)^7–9^. Based on the evidence from electroporation and electrofusion experiments, it has been suggested that the actin cortex is involved in the electroporation mechanism^10–12^. However, until now, a complete understanding of the role of the actin cortex in the electroporation of a lipid bilayer is lacking^13, 14^.

There have been several studies on the role of the cytoskeleton during electroporation of living cells^11, 12, 15–22^. In these studies, the actin filaments were manipulated by drugs that cause chemical disruption^11, 12, 15, 19^, stabilization of actin monomers,^16^ or by genetic engineering^17^ of the cytoskeleton. Particularly, it has been shown that actin depolymerization in Chinese hamster ovary (CHO) cells and human erythrocytes leads to acceleration of the post-pulse membrane resealing, while it does not considerably affect the initial steps in the permeabilization process^11, 12^. More recently, nanosecond pulses have gained attention due to their ability to permeabilize intracellular membranes^23^. Disruption of the actin cortex in CHO cells appears to make the cells more susceptible to nanosecond electroporation^18^. Also, nanosecond electroporation was found to be more damaging to non-adherent cultures such as Jurkat cells than adherent cell cultures such as HeLa cells, possibly because of the less extensive cytoskeletal network of the former^20^. Moreover, some biological processes like apoptosis and necrosis can be triggered by nanosecond pulses^19, 21^, which could involve the pulse-mediated disruption of the actin cytoskeleton^15, 22^.

The studies discussed above illustrate how challenging it is to decouple the mechanical interaction between actin filaments and the membrane from other possible biological processes involving actin, such as apoptosis and necrosis, and to manipulate the cytoskeleton of living cells without side effects like cell death. Therefore, simplified cell models like planar lipid bilayer models^24–26^ or giant unilamellar vesicles (GUVs)^27–29^ have been utilized to study the fundamental mechanisms of pore formation during electroporation. Studies on GUVs have revealed the presence of micrometer sized pores (referred to as macro-pores) during electroporation^7, 30–33^. The resealing time of these pores is typically in the order of 10 ms^30^, as opposed to several minutes in living cells. It has been shown that the dynamics of these pores can be modified by adjusting the edge tension of the lipid bilayer, e.g. by adding cholesterol to increase the edge tension^34, 35^. Moreover, gel-phase lipids have been shown to prevent the formation of macropores due to high surface viscosity^36, 37^. From these studies, it can be concluded that the membrane components are crucial for the dynamics of electropermeabilization in cells. However, GUVs and living cells exhibit drastically different permeabilization behaviors. The pores formed in GUVs due to pulse application can reach the micrometer range, and reseal within tens of milliseconds. By contrast, the electropores (or permeated membrane) in living cells were reported to be in the order of few nanometers^6, 38, 39^ and express a slow resealing process which takes minutes up to hours^38, 40, 41^, suggesting that the permeabilization dynamics are influenced by intracellular structures such as the actin cortex^10, 12^.

In this research, we focus on the role of the actin cortex during electroporation of the membrane in order to understand the discrepancy between the electropermeabilization of simplified model membranes and the plasma membrane. To isolate the mechanical role of the actin, we have incorporated a biomimetic actin cortex inside GUVs (See Figure 1A-B). We have used a high-speed imaging technique to reveal the pore formation and resealing in response to controlled electric pulses for GUVs, with and without an encapsulated actin network. Additionally, we assessed the permeability dynamics of both GUVs after electric pulses. Finally, we use confocal imaging to investigate the structural stability of the encapsulated biomimetic actin cortex in the electric field.

**Figure 1.**
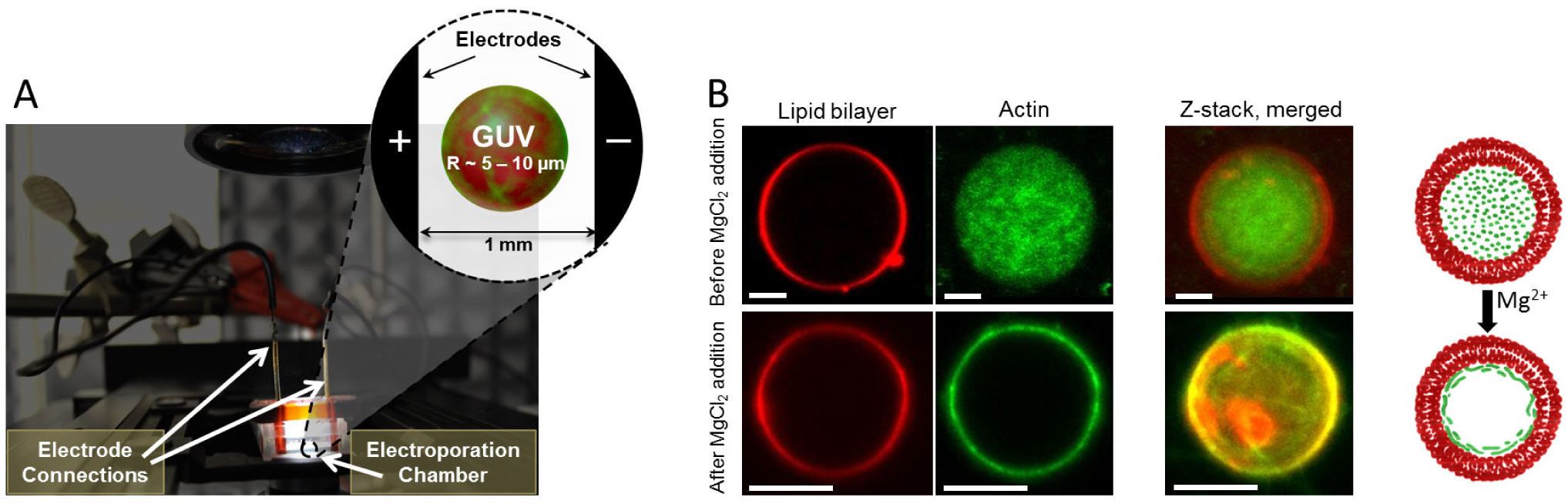
The electroporation setup. (A) Photograph of the electroporation setup, with a schematic of the GUV in between the electrodes shown in the inset, not drawn to scale. (B) confocal fluorescence microscopy images of a GUV (red signal, ex/em 560 nm/583 nm) before and after the addition of 19 mM MgCl_2_, showing that actin polymerizes and accumulates underneath the lipid bilayer membrane (green signal, ex/em 495 nm/519 nm). The scale bar in all images is 5 *µ*m.

## Materials and Methods

### Preparation of the GUVs

The GUVs (with and without actin filaments) were prepared by electroswelling method, based on a procedure utilized by Schäfer et al.^42^. The following membrane composition was used for both GUVs in the bright field experiments: 95 mol % 1,2-dioleoyl-*sn*-glycero-3-phospholine (DOPC) and 5 mol % Ca^2+^-ionophore (A23187). For the confocal experiments, the membrane was fluorescently labelled using the following membrane composition: 94.5 mol % DOPC, 5 mol % Ca^2+^-ionophore and 0.5 mol % 1,2-dioleoyl-*sn*-glycero-3-phosphoethanolamine-N-(lissamine rhodamine B sulfonyl) (ammonium salt) (DOPE-RhoB, ex/em 560 nm/583 nm). The lipids were purchased from Avanti Polar Lipids, Inc and the Ca^2+^-ionophore from Sigma Aldrich, and these compounds were stored at *–*20 °C. All membrane components were dissolved separately in chloroform at a concentration of 1 mg/ml, stored under nitrogen at *–*20 °C, and mixed prior to the experiments. 10 *µ*l of lipid mixture was deposited on the conductive face of two Indium Tin Oxide (ITO) slides (purchased from Sigma Aldrich) and left under vacuum for more than 2 hours. Afterwards, the ITO slides were inserted in a teflon swelling chamber with the conductive sides facing each other, spaced 1.5 mm apart. 335.6 *µ*l swelling buffer (pH of ∼8) was inserted in between the ITO slides, consisting of 200 mM sucrose, 2 mM Tris-HCl (Thermo Fischer), 0.17 mM Adenosine 5’-triphosphate disodium salt hydrate (Na_2_ATP) and 0.21 mM dithiothreitol (DTT) (both purchased from Sigma Aldrich). In the case of the GUVs with encapsulated filaments, monomeric actin from rabbit muscle (Thermo Fischer), purchased as 1 mg of lyophilised powder, was dissolved in 2 mM Tris-HCl (pH 8.0) to get a stock solution of 2 mg/ml and stored at *–*80 °C. For visualization of the filaments, fluorescently labeled actin monomers with Alexa Fluor 488 (Thermo Fischer, ex/em 495 nm/519 nm) was purchased as a solution with a concentration in the range of 3 *–* 5 mg/ml in a buffer (5 mM Tris (pH 8.1), 0.2 mM DTT, 0.25 mM CaCl_2_, and 0.2 mM ATP) containing 10% (w/v) sucrose. A solution of 7 *µ*M actin monomers, of which 0.5 *µ*M fluorescently labeled, was added to the swelling buffer. An alternating current (AC) was applied at 70 Hz, 2.4 V for 3 hours at room temperature (around 20 °C), using the Agilent 33220A 20 MHz Function / Arbitrary Waveform Generator.

### Polymerization and binding of actin to the membrane

After the swelling of the GUVs and encapsulation of 7.5 *µ*M actin monomers, actin polymerization at the membrane was initiated. For both the polymerization and the binding of the filaments to the membrane Mg^2+^ ions are used ions^42^, as shown in Figure 1B. The GUV solution obtained by electroswelling was diluted about 5 times with 1.3 ml glucose buffer of 200 mM glucose and 2mM Tris-HCl (pH of ∼8) in order to dilute the actin monomer concentration and the sucrose concentration on the outside of the GUVs. 163 *µ*l of 67.3 mM MgCl_2_ solution (pH of ∼8.5) was then added to increase the MgCl_2_ concentration to 6 mM. Subsequently, due to the presence of the Ca^2+^-ionophore in the membrane, the Mg^2+^ ions are transported into the GUVs to initiate polymerization and bind the actin filaments to the membrane as a cortex through electrostatic interactions^43^. Afterwards, the GUVs were left overnight at room temperature at the 6 mM MgCl_2_ and an 1.55 *µ*M actin concentration on the outside of the GUVs. Prior to the electroporation experiments, the GUVs were diluted ten times more with 1 ml of the 200 mM glucose buffer, in order to sediment the GUVs and enhance the optical contrast between the solutions inside and outside the GUVs. Afterwards, the MgCl_2_ concentration in the outer solution was increased to 19 mM to ensure the formation of a cortex at the membrane for all GUVs by the addition of 412.4 *µ*L of 67.3 mM MgCl_2_. In addition, this dilution further reduced the concentration of the actin outside the GUVs to 0.04 *µ*M. In our hands, this procedure led to the most consistent results, with a large majority of GUVs containing an actin network (*>* 90 %) (see Figure S.1 in the supplementary information). Since the ion concentration has a large impact on the response of the GUVs to electric pulses, altering the deformations of the GUVs^44^ and the pore dynamics^45^, we fixed the concentration of MgCl_2_ at 19 mM. The stability of the GUVs was not affected by the high ion concentration overnight. The absence of a detectable actin network on the outside of the GUV during the storage overnight and during the electroporation experiments was confirmed by the control experiments where empty GUVs are incubated with 1.55 uM and 0.04 *µ*M actin (see Figure S.2 in the supplementary information). Note that all of our data for electroporation were obtained at a dilute concentration of actin outside of the GUVs (C_*actin*_= 0.04 *µ*M), which is far below the critical concentration to form actin filaments^46^.

### Electroporation setup combined with high speed imaging

To image the dynamics of the membrane during an electric pulse, the GUVs were visualized by an inverted microscope (Zeiss Axio Observer.Z1) equipped with a Phantom v9.1 high-speed camera (10000 *–* 15000 frames per second, Vision Research Inc.) with a 40x oil immersion objective (Ph3, Plan-Neofluar, 40x/1.30), providing a pixel size of 0.3 *µ*m. Prior to the experiments, the imaging chamber was treated with 5 g/l Bovine Serum Albumin (BSA) solution (purchased from Sigma Aldrich) for 20 minutes, to avoid rupture of the GUVs upon contact with the glass bottom of the chamber. Custom made stainless-steel 20 mm-long electrodes with 1 mm distance were submerged in 1 ml glucose buffer with 412.4 *µ*l MgCl_2_ (with a concentration value of 67.3 mM) and 33 *µ*l of GUV solution was added by carefully pipetting the GUV-solution in between the electrodes using a cut-off pipette tip (Figure 1A). We selected the actin-encapsulated GUVs based on the fluorescence signal of the actin, to ensure that the filaments were organized along the GUV membrane as a cortex, as shown in Figure 1B. The actin-encapsulated GUVs had a radius between 5 and 10 *µ*m, empty GUVs of similar sizes were selected for the control experiments. During the dynamics experiments, consecutive 500 *µ*s pulses with increasing amplitude were applied. The electric pulses induce a transmembrane voltage as follows^47^:

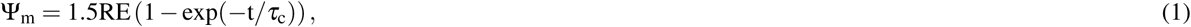

where R, E and t indicate the radius of the GUV, the applied electric field strength and the duration of the pulse, respectively. *τ*_c_ is the charging time of the membrane given by

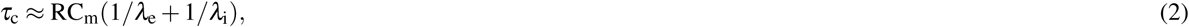

where C_m_, *λ*_e_ and *λ*_i_ are the membrane capacitance, and the internal and external solution conductivity, respectively. In our experiments the pulses were chosen such that the induced transmembrane voltage started from about 1 V, with approximately 1 minute interval between the pulses to minimize the effect of the former pulse. The sucrose-filled GUVs with glucose solution on the outside assure sufficient optical contrast to image the GUVs in bright field, and indicate permeabilization of the membrane by a contrast loss. Consecutive pulses with increasing amplitude were applied until a complete contrast loss of the GUVs was observed.

### Electroporation setup for pore dynamics

To assess the resealing of the membrane, the transport of sulforhodamine B through the membrane was monitored. Sulfurho-damine B was added to the outer solution at a concentration of 2.5 *µ*M. The same pulse protocol was used as during the high-speed imaging experiments. Dye transport was captured with a Zeiss LSM 710 inverted confocal microscope, using a 100x oil immersion objective (Ph2, Archoplan, 100x/1.25 Oil) and a sampling frequency of 1.5 Hz. To obtain the uptake kinetics of the sulforhodamine B, the intensity of 80 % of the inner GUV area was monitored, to exclude the increased intensity of the membrane. In addition, the intensity of the background was captured. The data was normalized as follows: I_uptake_ = (I_dye,t_ *−* I_dye,0_)/(I_background,t_ *−* I_dye,0_), where I_uptake_, I_dye,t_, I_dye,0_ and I_background,t_ represent the normalized intensity inside the GUV at time *t*, the measured intensity inside the GUV at time *t*, the initial intensity prior to the pulse and the intensity of the background, respectively.

### Electroporation chamber combined with confocal imaging

The response of the actin cortex to the electric pulses was visualized with a Zeiss LSM 710 inverted confocal microscope, using a 100x oil immersion objective (Ph2, Archoplan, 100x/1.25 Oil) to image both the lipid bilayer and the actin cortex. Two different pulsing protocols were used: (i) consecutive increasing pulses starting from low fields, increasing gradually to higher fields (ranging from 10 *–* 300 V/mm with steps of approximately 10 V/mm), (ii) immediate high pulses (ranging from 300 *–* 1000 V/mm with steps of approximately 200 V/mm), where only 2 to 4 pulses were applied of increasing field. In both cases 500 *µ*s pulses were used, separated by approximately 1 minute. The first pulse protocol (i) was used to test the onset of the breakdown of the actin cortex. The second pulse protocol (ii) was used to decouple the breakdown of the actin cortex from the cumulative effect of the multiple consecutive pulses. A z-stack of 7 focal planes, with a range of approximately the diameter of the GUV (Δz ∼ 4 *µ*m), of the fluorescence signal of the membrane and the actin was taken before and after the pulses. The z-stack imaging is a slow technique (approximately 5 to 10 seconds time interval in between the z-stacks), so only the GUV size and the actin cortex stability before and after the pulse were extracted.

### Data analysis of GUVs response

The bright field images of the GUVs were analyzed by a custom written Matlab script. The contours of the GUVs were detected with the Canny edge detection method and fitted to an ellipse to extract the equatorial (a) and polar (b, the distance from the center to the poles of a spheroid) radii of the GUVs. For the dynamics studies, these two radii were used to obtain the deformation ratio (a/b) during and after the pulse. Afterwards, the time-dependent deformation was fitted to an exponential decay curve to obtain the relaxation time(s) of the GUVs^30^ (Figure 2).

**Figure 2.**
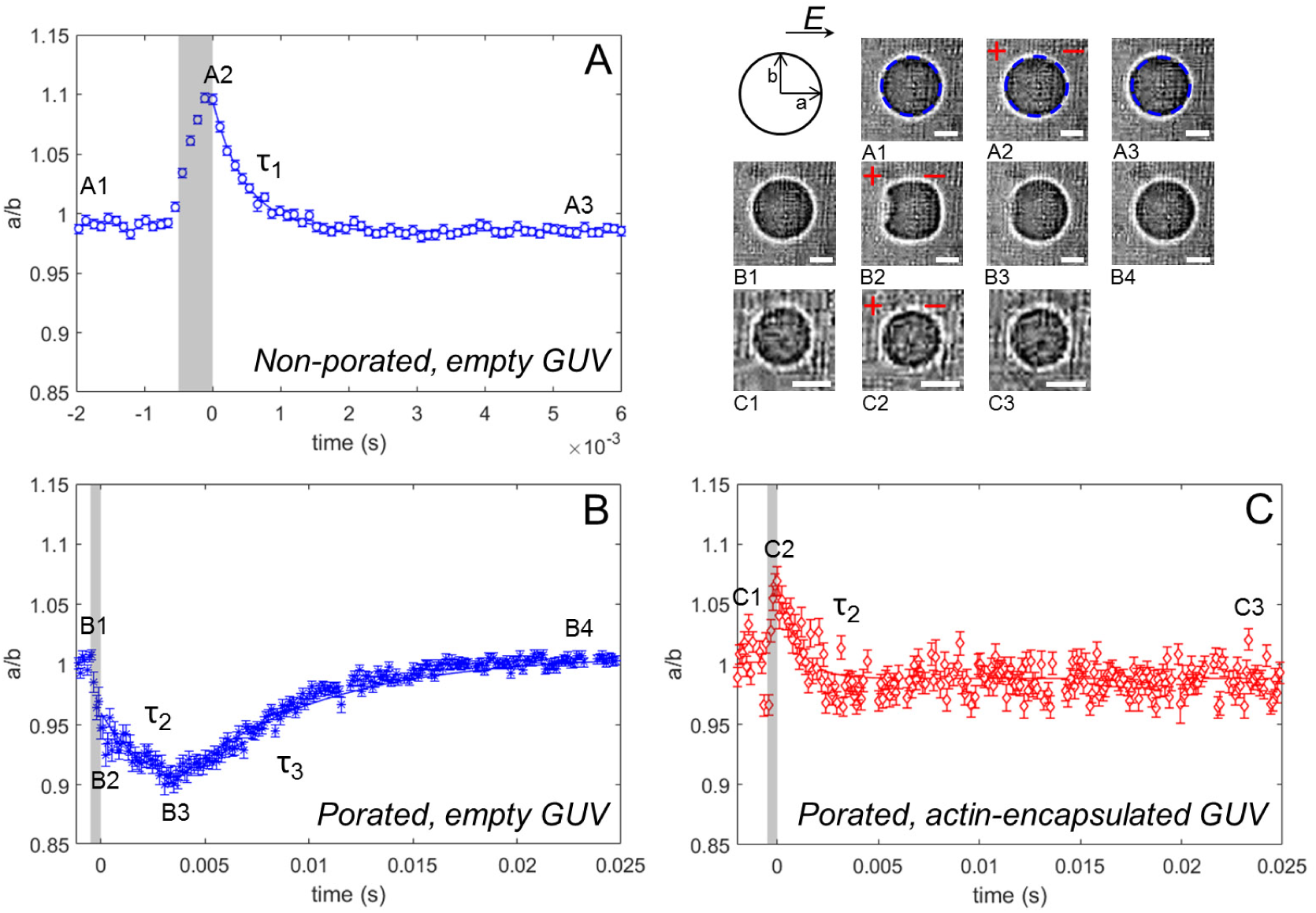
The relaxation dynamics of GUVs with and without an encapsulated actin shell. (A) The dynamics of a non-porated empty GUV, where deformation and a single relaxation (*τ*_1_) is observed. The blue dotted lines in the snap shots of A1-A3 show the results of the tracking method used to determine radii a and b. (B) The dynamics of a porated empty GUV, where a pore is observed together with a double relaxation (*τ*_2_ and *τ*_3_). (C) The dynamics of a porated actin-encapsulated GUV where no macropore is observed, and only a single relaxation (*τ*_2_).The relaxation times are obtained from exponential fits to the deformation data, as explained in the text. The scale bar in all images is 5 *µ*m. The definitions of non-porated and porated GUVs are explained in the text.

During the confocal experiments, multiple planes of the GUVs were imaged. The radius of the GUVs was determined from the fluorescent confocal membrane images taken at the equatorial plane of the GUV by a similar detection method as described above. These images were used to obtain the area loss of the GUVs as a function of electric field. In order to remove spurious effects due to motion of the GUV in and out of the imaging plane (which could change their observed radius), the normalized area (A_norm_) was calculated at pulse *n* and corrected for the area loss at the previous pulse (*n −* 1): A_norm_ = (A_f_/A_i_)_n_(A_f_/A_i_)_n*−*1_. Here, A_i_ and A_f_ represent the initial and final area of the GUV before (initial) and after (final) the pulse, respectively. (A_f_/A_i_)_n_ and (A_f_/A_i_)_n*−*1_ are the normalized areas of the GUV at the pulse *n* and at the preceding pulse (*n −* 1). By multiplying these two factors, the normalized area is corrected for shrinkage due to earlier pulses. Additionally, the intensity of the fluorescence signal of the actin was obtained from confocal z-stacks by determining the mean and average intensity of all planes. Photobleaching of the actin cortex was removed by correcting the data at scan number *k* with a reference of a separate photobleaching experiment of a GUV without applying any pulses. By normalizing the actin fluorescence intensity from the bleaching experiment, the correction factor for the photobleaching per scan is calculated (I_ref,k_ = I_k,ref_/I_0,ref_, where I_0,ref_ and I_k,ref_ are the intensities of the image at the start of the photobleaching experiment and at the relevant scan number *k*, respectively). Consequently, the results of the electroporation experiments were corrected as follows: I_norm_ = I_k_/(I_0_I_ref,k_). Again, I_0_ and I_k_ represent the intensities of the image at the start of the experiment and at scan number k, respectively, in this case during an electroporation experiment (see Figure S.3 in the supplementary material).

## Results

### The dynamics of the GUVs

To investigate the influence of the actin cortex on the electroporation of GUVs, we captured the dynamics of the GUVs by high speed imaging (10000 *–* 15000 fps) in bright field. The deformations of the GUVs were measured in terms of the deformation ratio a/b during and after the electric pulse (see the schematic in Figure 2). The GUVs are electroporated once the accumulated transmembrane voltage exceeds a threshold voltage. Electroporation can be recognized by the observation of a macropore or by the contrast loss due to the exchange of glucose and sucrose molecules between the interior and exterior of the GUV. Below the critical threshold field, where no macropores nor contrast loss were observed, the GUVs exhibited squarelike deformations, as shown in Figure 2A. These deformations relaxed back exponentially with a characteristic relaxation time (*τ*_1_) in the order of 100 *µ*s for both the empty and the actin-encapsulated GUVs. Riske and Dimova^44^ previously showed that the presence of the ions (NaCl, Ca^2+^ or Mg^2+^ acetates) flattens the membrane and causes squarelike deformations, which they attributed mainly to the electrophoretic forces of the ions.

The conductivity ratio *χ* is defined as the ratio between the conductivity of the internal (*λ*_in_) and external solution of the vesicle (*λ*_out_), *χ* = *λ*_in_*/λ*_out_. Depending on this conductivity ratio, vesicles are expected to either deform along the field, referred to as a tubelike shape, (deformation ratio a/b *>* 1, *χ >* 1) or perpendicular to the field, referred to as a disklike shape, (deformation ratio a/b *<* 1, *χ <* 1). We observed both disklike and tubelike deformations during the pulsing experiments of both the empty and the actin-encapsulated GUVs. It must be noted that an ionophore is used to transport Mg^2+^ into the GUVs to polymerize actin and construct the biomimetic cortex after the GUV formation, as discussed in the methods section. Some variation in the conductivity of the inner solution of the GUVs is possible due to our preparation method, which would explain the observations of both disklike and tubelike deformations. Additionally, permeabilizing pulses cause an exchange of the inner and outer solutions, and consequently may alter the conductivities of the two solutions. As these observations are similar for both the empty and actin-encapsulated GUVs, they can be related to the presence of the ionophore and the ion-imbalance of the inner and outer solution.

The response of the empty and the actin-encapsulated GUVs to an electroporative pulse was strikingly different, as shown in Figure 2. Macropores were observed for the empty GUVs, as was reported before^30^, whereas the actin-encapsulated GUVs exhibited no visible macropores. Additionally, for the empty GUVs we observed a characteristic relaxation process, where the relaxation of the macropore (with a duration *τ*_2_ of ∼1 ms) was often followed by another relaxation event (with a duration *τ*_3_ of ∼ 10 ms), as illustrated in Figure 2B and movie S1. This response agrees with previous studies on empty GUVs composed of egg phosphatidylcholine by Riske and Dimova^30^. They attributed *τ*_2_ to the relaxation time of the macropore in a standalone lipid bilayer (*τ*_pore_ ∼ *η*_s_r/(2*γ*), where *η*_s_, *γ* and r are the surface viscosity, the line energy per unit length and the pore radius, respectively) and *τ*_3_ to the relaxation of the membrane due to either the excess surface area of the GUVs or an increase of the excess area during the macroporation^30^. The similarities between the macropores exhibited by our empty GUVs and previous studies show that the ionophore and the Mg^2+^-ions present in the solution have a negligible effect on the GUV dynamics during electroporation. In a clear contrast to empty GUVs, the actin-encapsulated GUVs showed only small deformations during the electroporative pulses and did not exhibit macropores (Figure 2C and movie S2). Since no macropore was observed for these GUVs, electropermeabilization was determined by the contrast loss of the GUVs. This contrast loss indicates the permeabilization of the membrane, due to the exchange of molecules through the membrane. Due to limited recording time of the high-speed imaging in the experiments (maximum of 2 seconds), no long-term dynamics of the contrast loss could be captured.

The relaxation time *τ*_2_, which characterizes the time-dependent deformation of the actin-encapsulated GUVs in the electroporative regime, is comparable to the relaxation time *τ*_2_ of the empty GUVs (Figure 3A). However, the slow dynamics of the contrast loss after a pulse for the actin-containing GUVs indicates that the membrane remains permeabilized up to minutes, which was not observed for the empty GUVs. Therefore, the pore resealing time of the actin-encapsulated GUVs cannot be deduced from *τ*_2_. Previous studies on living cells have proposed that the formed electropores in the membrane cannot expand beyond the mesh size of the actin network^13, 38^. Possibly, a similar mechanism limits the growth of the pores in the actin-encapsulated GUVs.

**Figure 3.**
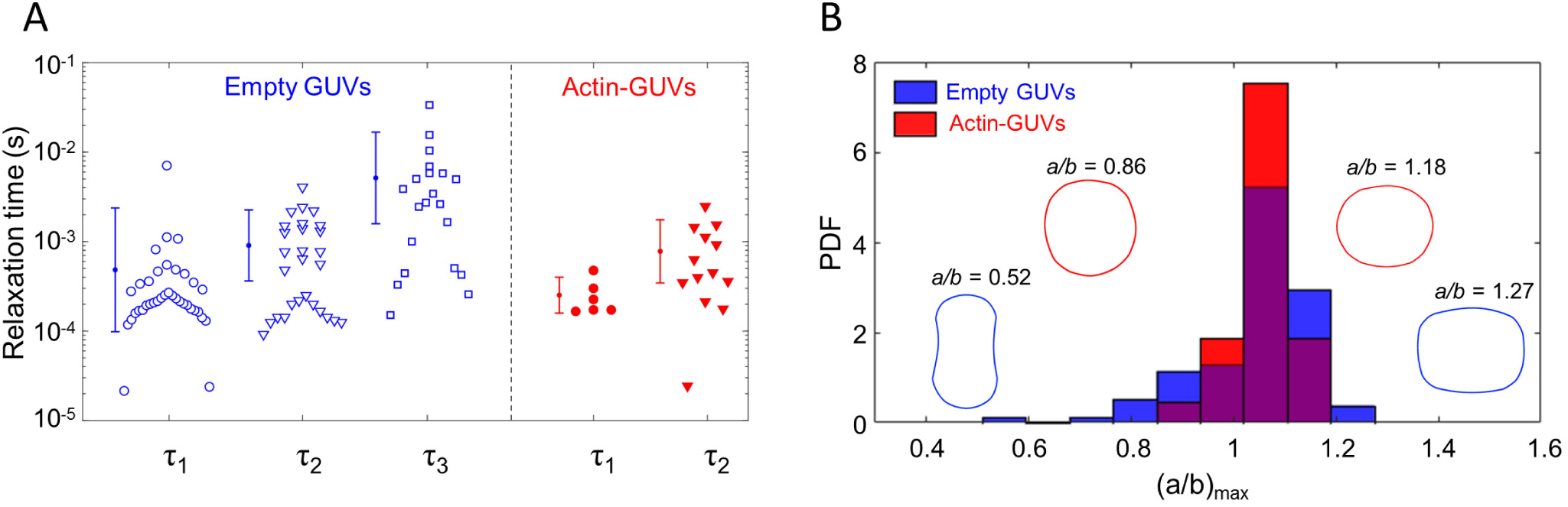
The relaxation time and maximum deformation of the GUVs during an electric pulse with and without actin shell. (A) Relaxation times of empty (blue open symbols) and actin-encapsulated (red filled symbols) GUVs in the non-electroporative regime (*τ*_1_) and the electroporative regime (*τ*_2_ and *τ*_3_). (B) The distribution of the maximum deformations of empty (blue) and actin-encapsulated (red) GUVs during all pulses (both electroporative and non-electroporative). The schematics in the histogram represent simplified contours of the corresponding disklike and tubelike deformations, not to scale. The data in both panels is of 16 actin-encapsulated GUVs with an average radius of ∼5 *µ*m (ranged from 3 to 6.4 *µ*m) and 10 empty GUVs with an average radius of ∼5 *µ*m (ranged from 3 to 6.7 *µ*m). In both cases the experiments have been repeated five times on different days.

As discussed above, we observed both disklike (a/b *<* 1) and tubelike (a/b *>* 1) deformations for the empty and the actin-encapsulated GUVs. For both types of GUVs, as shown in Figure 3B, the majority of vesicles undergoes a tubelike deformation, indicating a higher conductivity of the interior of the GUVs due to the Mg^2+^ transport across the membrane by the ionophore. The deformations of the actin-encapsulated GUVs are significantly smaller than those of the empty GUVs, which can be seen from the more narrow distribution of (a/b)_max_ (shown in red in Figure 3B). Experiments on GUVs in alternating current (AC) fields have shown that the extent of membrane deformation depends on the bending rigidity of the membrane^48^. Additionally, in DC field experiments, non-electroporative pulses can be used to study the stiffness of the membrane^49^. In our experiments we focused on pulses around the transmembrane voltage of 1V, therefore we cannot derive the precise mechanical properties. Nevertheless, the smaller deformations of the GUVs with an encapsulated actin shell clearly signify an increase in the bending rigidity of the GUVs, consistent with previous studies employing either AFM-indentation, micropipette aspiration, or flicker spectroscopy^42, 50, 51^. The wide distribution observed for (a/b)_max_ of the GUVs is caused by the preparation technique. The electroswelling method unfortunately offers a poor control on the membrane tension of the GUVs (both for the empty and the actin-encapsulated GUVs). Moreover, the actin-encapsulated GUVs likely possess different actin shell thicknesses (see Figure S5 in the supplementary material), leading to different mechanical properties^42^. In conclusion, our results show that the actin cortex prevents macropore formation and reduces membrane deformation, which may contribute to the differences observed between the response of cells and empty GUVs to electric pulses.

### Resealing of the permeabilized membrane

The permeabilization dynamics of the membranes has been determined by the uptake of sulforhodamine B molecules into the GUVs. Due to the relatively slow imaging, only the transport after the pulse could be captured, which is predominantly diffusive^52^. The applied pulses are in the same range where macro-pores were observed for the empty GUVs. An increase in the inner fluorescence of the GUVs immediately after the pulse is defined as the point where electroporation occurs. To find the amount and the characteristic time of dye uptake after the pulse, we use the following fitting equation^52^:

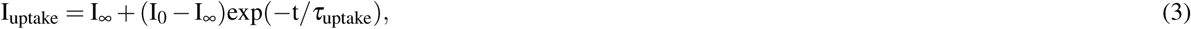

where I_∞_, I_0_ and *τ*_uptake_ represent the final sulforhodamine B intensity of the GUV, the initial sulforhodamine B intensity of the GUV prior to the pulse and the characteristic time of the uptake of the dye molecules. The two fitting parameters I_∞_ and *τ*_uptake_ for the actin-encapsulated GUVs after a pulse are larger than for the empty GUVs (Figure 4). A former study on the dye leakage of GUVs employing a high temporal resolution showed that the leakage of dye mainly occurred through macropores^53^. Similarly, we observed macropore formation for the empty GUVs, which was accompanied with ∼2 % uptake of dye molecules (∼0.02 I_∞_) after a single pulse and a characteristic uptake time of approximately 12 seconds. Taken together, these observations indicate that most dye transport occurs during the pulse and shortly after the pulse. It must be noted that earlier studies have shown a resealing of the membrane in 10 ms^30^, which is faster than our temporal resolution. Therefore, the actual characteristic uptake time might be considerably smaller than 12 seconds, although the presence of the ionophore in the membrane may slow down membrane resealing. The actin-encapsulated GUVs, on the other hand, do not show any macro-pores and exhibit a longer characteristic uptake time of approximately 146 seconds together with a dye uptake of ∼38 %. Our observations are reminiscent of prior observations of a high and long-lived permeability for lipid membranes associated with an agarose mesh work^53^. In that work it was proposed that the agarose mesh stabilizes pores formed by an electric pulse. The actin network in our experiments appears to affect the membrane stability in a similar fashion, enabling transport through the membrane for a longer duration than for the empty GUVs.

**Figure 4.**
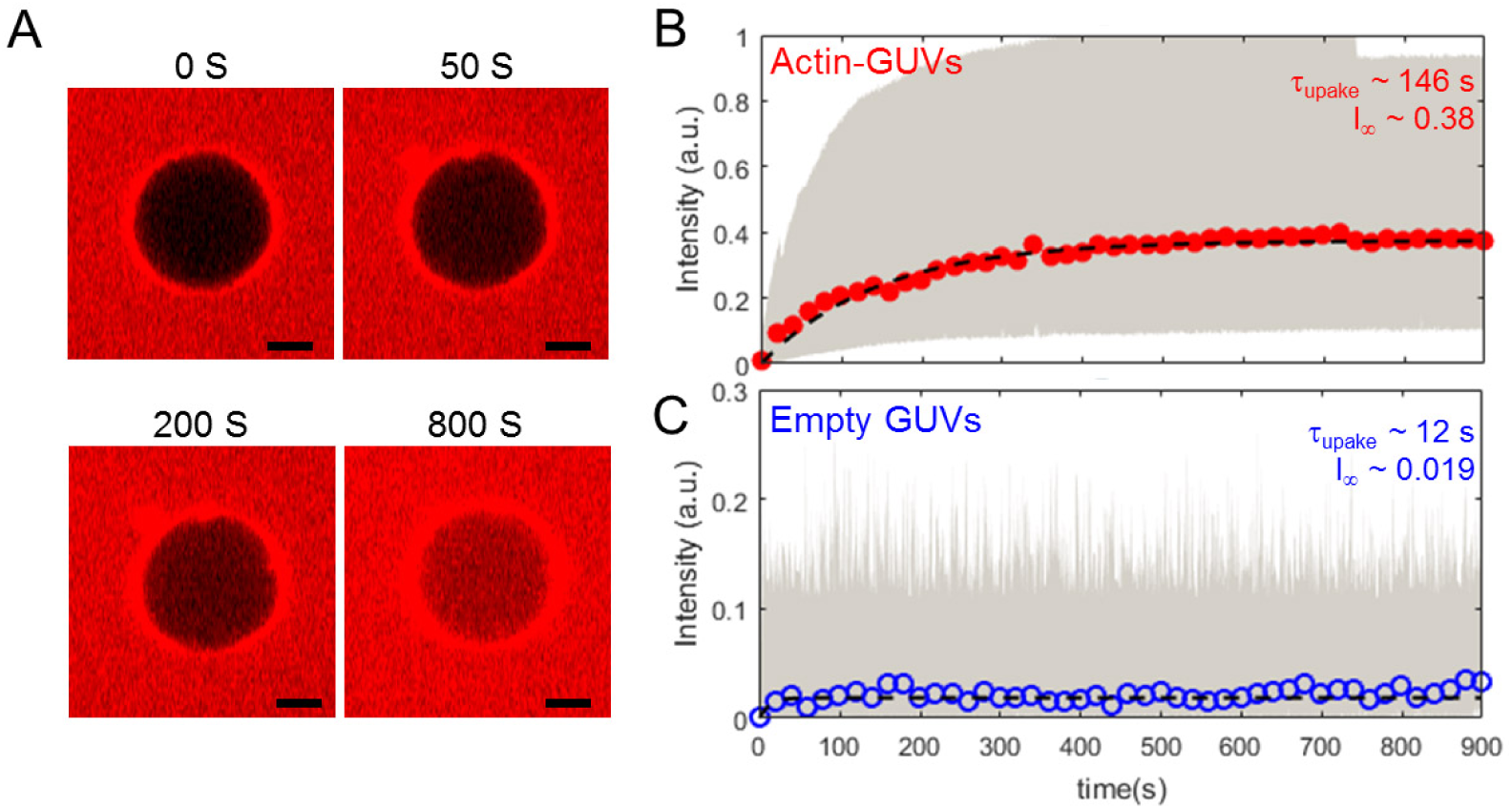
The kinetics of dye uptake during the resealing of GUVs. (A) Four snapshots of a resealing experiment of an actin-encapsulated GUV, showing the uptake of dye molecules from the outside solution after a 85 V/mm pulse over time. The scale bar in all images is 5 *µ*m. (B) The average intensity of sulforhodamine B in the actin-GUVs over time of 13 different GUVs with an average radius of ∼10 *µ*m (ranged from 5 to 17 *µ*m). The GUVs were exposed to electroporative pulses inducing a transmembrane voltage above 1 V. Already one single 500 *µ*s pulse caused considerable uptake of the dye molecules. (C) The average intensity of fluorescent dye in empty GUVs over time of 15 different GUVs. To ensure dye uptake by the GUVs after pulse application, multiple pulses were applied when no visible dye uptake was obtained. Only the pulses where an increase of the dye intensity immediately after the pulse was observed have been selected to calculate the average intensity increase. The dotted lines in both graphs represent the least-squares fit of Equation 3 through the averaged data. The highlighted area in gray in both graphs shows the spread of the data points of all experiments. Only every twentieth data point of the averaged data is shown here, to improve the readability of the graph. The fitting parameters, characteristic time and amount of uptake of dye molecules, obtained from the fits are shown in the graphs.

Despite the difference between the structure and organization of our biomimetic cortex and the actin cortex of a living cells, similarities can be found between their response to electric pulses. The exponential increase in dye uptake of the actin-encapsulated GUVs appears similar to the exponential uptake of living cells^52^. The presence of an actin network is known to increase the membrane tension^54, 55^. The enhanced tension likely explains the slow resealing process of permeabilized structures in actin-encapsulated GUVs, compared to empty ones. A slow resealing process of the membrane is consistent with observations in electroporated cells, where the actin cortex influences the resealing of the permeated structures^11, 12^.

### Stability of the actin network

To assess the response of the actin network inside the GUV to electric pulses, the membrane and actin were concurrently visualized by confocal microscopy. The fluorescence signals of the membrane and actin network were used to detect the area of the GUV and the actin intensity, respectively (see Figure S.4 in the supplementary material). In order to determine the coverage of the actin network on the inner membrane surface and any possible lateral inhomogeneities of the shell, a confocal z-stack was taken before and after the pulse. Consecutive pulses of increasing voltage were applied to the GUVs, with at least 1 minute in between the pulses to minimize the effect of the former pulse. Similar to earlier studies^7, 31, 34, 56^, the empty GUVs exhibit increasing shrinkage with higher electric pulses (Figure 5A). Strikingly, a single pulse of 300 V/mm does not induce the same shrinkage as a 300 V/mm pulse preceded by multiple pulses at lower field strength (see Figure 5A). Thus, multiple pulses exert a cumulative effect on GUV shrinkage. Compared to the empty GUVs, the actin-encapsulated GUVs display shrinkage at higher electric fields, i.e. the actin-supported bilayer has a higher electrical stability (Figure 5B). This could be caused by the increased surface viscosity of the bilayer due to the presence of the actin layer^37^.

**Figure 5.**
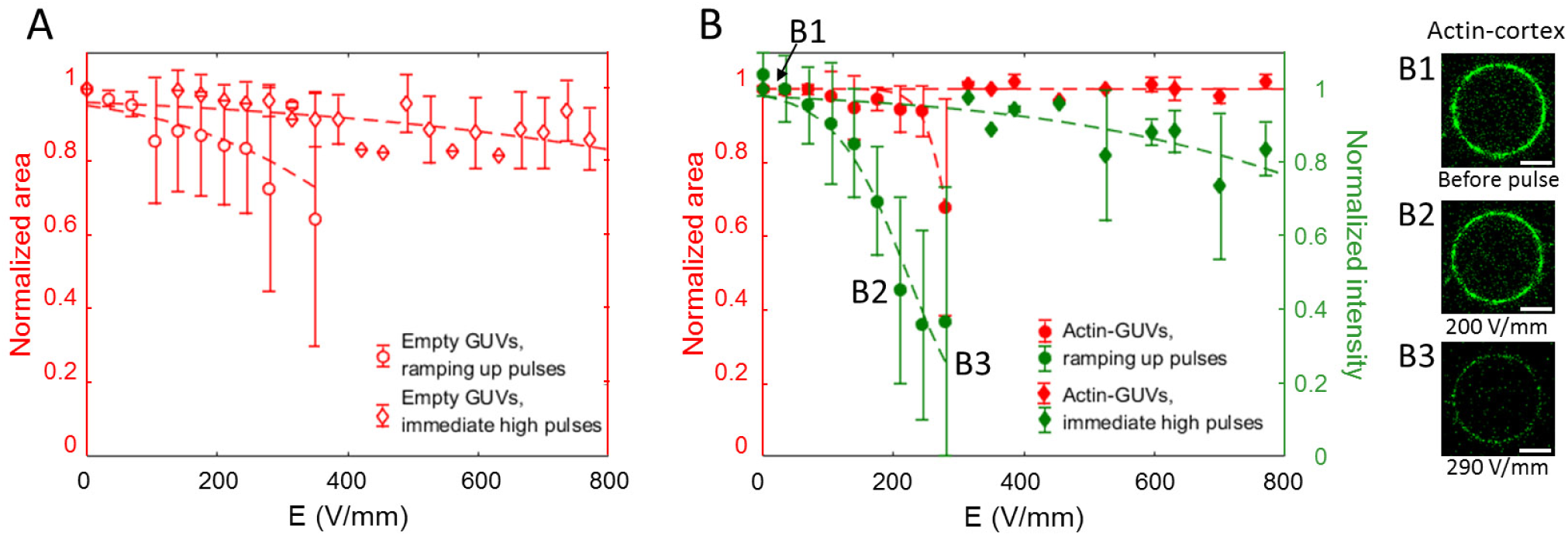
The structural stability of empty and actin-encapsulated GUVs. (A) The area loss of empty GUVs as a function of electric field. The average normalized area is shown for 15 empty GUVs in total with an average radius of ∼10 *µ*m, of which 9 have been exposed to the consecutive ramping up pulses and 5 to immediate high pulses. These experiments have been repeated four times on different days. (B) The area loss of the actin-encapsulated GUVs as a function of the electric field, which is associated with an intensity loss of the actin shell. The average normalized area is shown of 17 actin-encapsulated GUVs in total with an average radius of ∼10 *µ*m, of which 8 have been exposed to the consecutive ramping up pulses and 9 to immediate high pulses. These experiments have been repeated five times on different days. The confocal images show solely the actin fluorescence signal where the intensity loss can be attributed to shell disruption (B1, B2 and B3). Photobleaching of the fluorescence intensity of the actin shell is observed (shown in Figure S.3 in the supplementary information) and is corrected as discussed in the Materials and Methods section. The dotted lines in the graphs represent a least-squares fit of a sigmoid curve to guide the eye. For more statistics on the response of the GUVs, see Section S10 in the supplementary material. The scale bar in all images is 5 *µ*m.

Strikingly, the increasing pulses also result in a decreasing fluorescence signal of the actin at ∼150 V/mm, followed by GUV shrinkage at higher electric pulses (∼200 V/mm), as illustrated in Figure 5B and Movie S3. We observe that the decrease in the fluorescence signal of the actin is associated with a breakdown of the actin network. Additionally, the intensity loss of the actin layer due to the electric pulses appears to be slightly more at the poles, where the GUV is facing the electrodes (Movie S3). However, this radial dependence is not always observed clearly, possibly due to GUV’s rotation. After the breakdown of the actin network we did not observe any increase in the intensity in the center of the GUV (see Section S.5 and Figure S.4 in the supplementary material). By recording the photobleaching of the actin signal in the absence of the pulse and also the response of GUVs with encapsulated actin network to immediate high pulses, we could confirm that the fluorescence decrease is caused by the breakdown of the network and is not a side effect of bleaching. This notion is further supported by the observation that after the pulse, the actin fluorescence signal progressively decreased for several tens of seconds. This gradual decrease in the signal indicates a slow breakdown of the actin network. In addition, the shrinkage of the actin-encapsulated GUVs at higher electric fields, compared to empty ones, indicates that initially the actin network stabilizes the membrane and that shrinkage only sets in after the breakdown of the network.

Our observations are reminiscent of the depolymerization of actin crotex that has been observed for human cell lines^15^ and plant cells^16^ exposed to nanosecond pulsed electric fields (nsPEF). Different mechanisms have been proposed to account for actin depolymerization, including a direct effect of nanosecond pulses^15, 16^, osmotic swelling^22^ or biological processes^57^. In the GUV model system, we can exclude biochemical processes and in addition no osmotic swelling was observed in our experiments. The only factors present that might cause actin depolymerization are the mechanical forces, originating from the induced electrical membrane stress during the pulse, as well as the electrophoretic forces acting on the actin filaments as soon as the membrane is permeabilized.

We therefore estimate the magnitude of both the mechanical and the electrophoretic forces, considering the actin filaments as semiflexible polymers. For simplicity, we ignore the presence of actin bundles (Figure S.1 in the supplementary information). Assuming a uniform distribution of actin filaments over the membrane surface, the total number of filaments underneath the lipid bilayer can be estimated^58^ as:

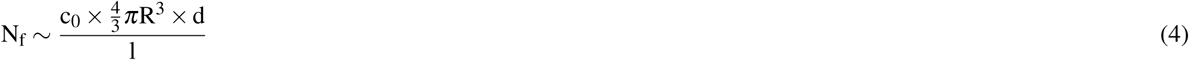

where c_0_ and d are the initial concentration (4.2 × 10^21^ m^*−*3^) and size of actin monomers, and l is the average length of the filaments. Assuming the filaments have a length in the order of l ∼ 4*µ*m (see Figure S.1 in the supplementary material) and with R = 10 *µ*m and d = 2.5 nm, we estimate the total number of filaments per GUV to be N_f_ ∼ 10^4^. The mesh size of the filament network can be approximated as: 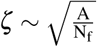 with A being the surface area of the membrane. The amount of stretch imposed on the length of connections in the actin network reads:

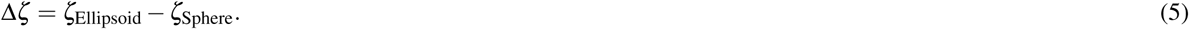

As the total enclosed volume of the GUVs is conserved during their deformation into ellipsoidal topologies in the experiments, the equatorial and polar radii become b = e^*−*1/3^R and a = e^2/3^R, respectively, where 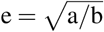 is taken from the example GUV shown in the schematic in Figure 2E. We choose a value of e ∼1.18, corresponding to the maximum deformation experienced by the GUVs. The total surface area of the GUVs increases during the deformation and hence the concomitant mesh size stretches by Δ*ζ* ∼ 0.8 nm (if we assume affine deformations for the network). Such an increase in the length of interconnected filaments induces a maximum mechanical force of the order of f_m_ ∼ 34 pN for a stretching stiffness of ∼48 pN/nm^59, 60^. This value is markedly smaller than the force needed for either initiating the depolymerization of a filament network^61^ or the rupture of single filaments^62^, which is in the range of ∼100 − 400 pN. It is, therefore, unlikely that the mechanical forces generated by in-plane tensions are the only origin of the breakdown of the actin network in our experiments. Other mechanisms, including electrophoretic forces, are expected to be involved.

As soon as the membrane is permeabilized by the electric fields, the membrane tension can relax back through the expansion of the pores and the release of the interior fluid. Additionally, upon applying an electric field on any charged molecules in a bulk solution, they experience a driving force. This effective force can drive and direct the motion of a free filament in the bulk fluid^63^. In contrast, when entangled and hindered from motion in a network (which is the case for our actin shell), the filaments can undergo mechanical forces between their building monomers. The force acting on the filaments in the shell due to the electric field is defined as^63^:

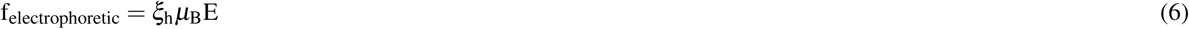

where *ξ*_h_ and *µ*_B_ represent the hydrodynamic friction coefficient per unit length of a filament close to the surface and the electrophoretic mobility of the actin measured in bulk solution, respectively. By assuming an average length of the actin filaments of ∼4 *µ*m and an interaction of the complete filament with the membrane, due to Mg^2+^-mediated adhesion, the force per unit length can be converted in the electrophoretic force f_electrophoretic_. The maximum force experienced by the actin filaments corresponds to a condition in which the filaments are perpendicular to the field. As soon as the GUV is permeabilized and pores are formed, the electric field penetrates into the GUV, with a maximum estimated value of ∼0.8E at the poles where the GUV is facing the electrodes (see Figure S.7 in the supplementary material). The fluorescence signal of the actin network drops at approximately 150 V/mm (Figure 5A). At this electric field and considering a hydrodynamic friction of *ζ* = 0.034 N.s/m^2^ (for cytoplasmic fluid motion perpendicular to the filament length^64^) and an electrophoretic mobility of *µ*_*B*_ = 10^*−*8^ m^2^/(V.s), we predict an electrophoretic force of f_electrophoretic_ ∼160 pN acting on a single filament for a vesicle size of 10 *µm*. Compared to the mechanical forces calculated above, these forces appear most plausible to initiate the disruption of the actin network. Importantly, the generated heat due to joule heating is estimated to be small (less than 3 K) in our experiments (see Section S9 in the supplementary material). Moreover, the disruption of the network mostly occurs above the critical transmembrane voltage (see Figure S.8 in the supplementary material), enabling the electrophoretic forces to affect the actin filaments.

## Conclusion

To shed light on the role of the actin cortex on the electroporation of cells, we have prepared GUVs with an encapsulated actin network that forms a biomimetic cortex and exposed them to electric pulses. Time-lapse imaging of the membrane revealed that membrane rigidification by the actin network^42^ inhibits large deformation of the GUVs during the electric pulses. Additionally, we found that the actin layer prevents the formation of large pores, referred to as macropores. Finally, we observed that membrane resealing after pulse application takes significantly longer in the presence of the actin shell than for a bare membrane. Interestingly, time-lapse imaging of actin revealed that, at higher electric fields, the electric pulses cause depolymerization of the actin shell. Based on the estimation of the relevant forces, we suggest that actin shell disruption is predominantly triggered by the electrophoretic forces on the actin filaments during the pulses.

Although our biomimetic cortex is still far from mimicking the actual actin cortex of a living cell, the behavior of the actin-encapsulated GUVs already shows the relevance of modeling the plasma membrane more closely. Our results provide the first step towards understanding the major differences between the electroporation of living cells and GUVs such as (lack of) macropore formation and the resealing dynamics of the defects. Consequently, these actin-supported GUVs enable exploring more complex processes, such as the mechanism of electro-gene transfection.

## Acknowledgments

DLP, AV, LR, and PEB thank the European Research Council (ERC) under the European Union’s Seventh Framework Programme (FP/2007-2013), grant agreement No 337820 (E-DNA-T-PEP). YM and GHK were supported by the European Research Council (ERC) from an ERC Starting Grant (335672-MINICELL). The authors thank Shaurya Sachdev for critical reading of the manuscript.

## Author contributions statement

P.E.B and D.L.P designed the experiments, D.L.P, V.K. and L.S. conducted the experiments. A.V. and L.R did the theoretical analysis. P.E.B., D.L.P., A.V., M.T.K., Y.M., A.M. and G.H.K contributed to the analysis and the interpretation of the data. D.L.P. and A.V. wrote the manuscript. P.E.B., D.L.P., A.V., L.R., M.T.K., Y.M. and G.H.K reviewed the manuscript.

### Additional information

#### Supplementary information

accompanies this paper at http://www.nature.com/srep

#### Competing interests

The authors declare no competing interests.

